## Supplementary Figures and Tables for "Chromatin-accessibility estimation from single-cell ATAC data with scOpen"

May 10, 2021

### Supplementary Figures

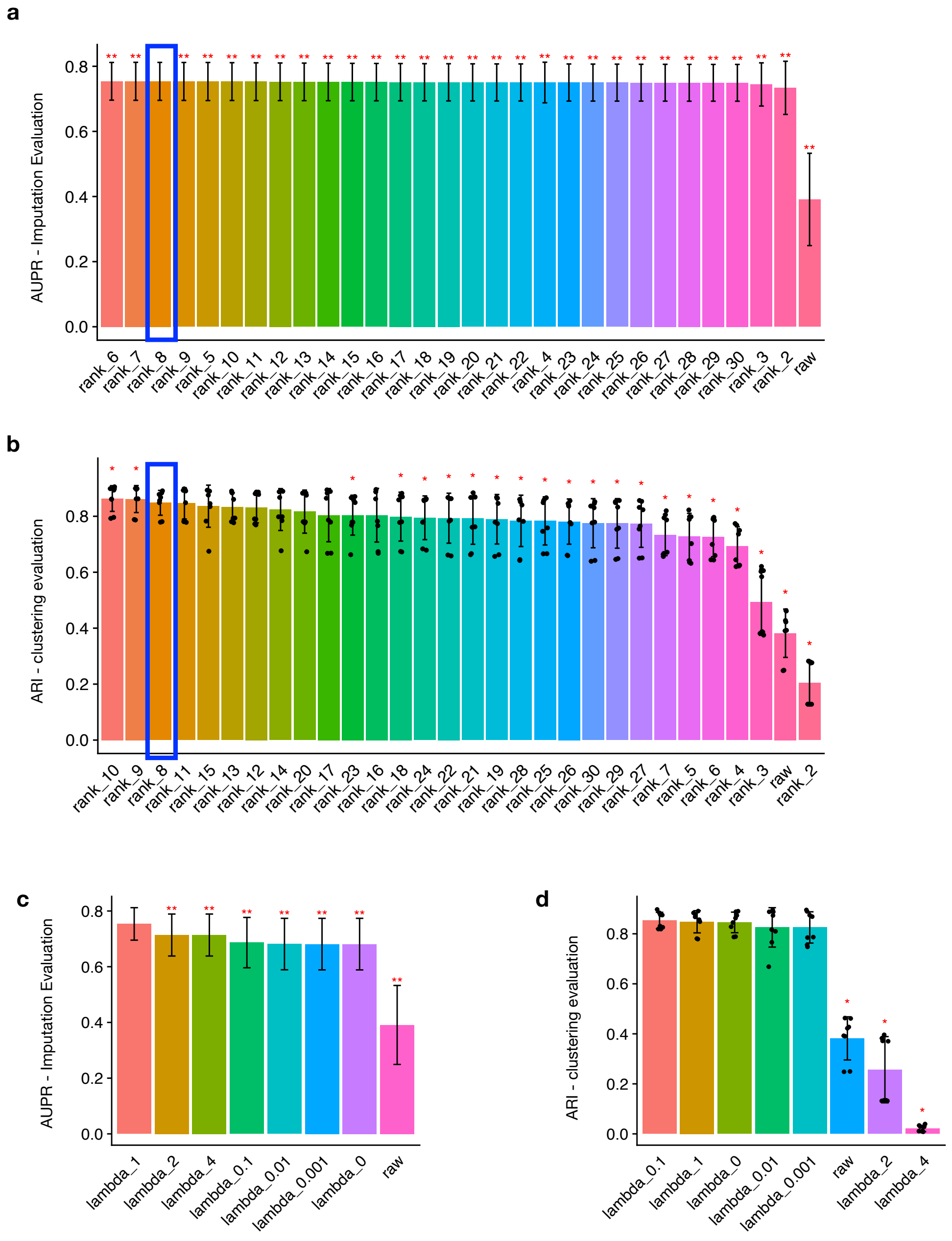


**Supplementary Fig. 1. Evaluation of hyper-parameters in scOpen using simulated dataset. a,** Bar plot showing mean AUPR of cells (imputation evaluation) by using different ranks in simulated dataset. Rank with 8 was selected by scOpen (highlighted with blue box). Error bars represent standard deviation. Asterisks denote statistical significance by comparing rank=8 to other values (** $p < 0.01$(Wilcoxon Rank Sum test, two-tailed, corrected using BH method for multiple comparisons). **b,** Same as **a** for ARI metric (clustering evaluation). **c,** Bar plot showing mean AUPR of cells (imputation evaluation) using different $\lambda$ in simulated dataset. Error bars represent standard deviation. We compared between $\lambda = 1$ and other values (** $p < 0.01$ and * $p < 0.05$ with Wilcoxon Rank Sum test, two-tailed, corrected using BH method for multiple comparisons).**d,** Same as **c** for ARI metric (clustering evaluation).

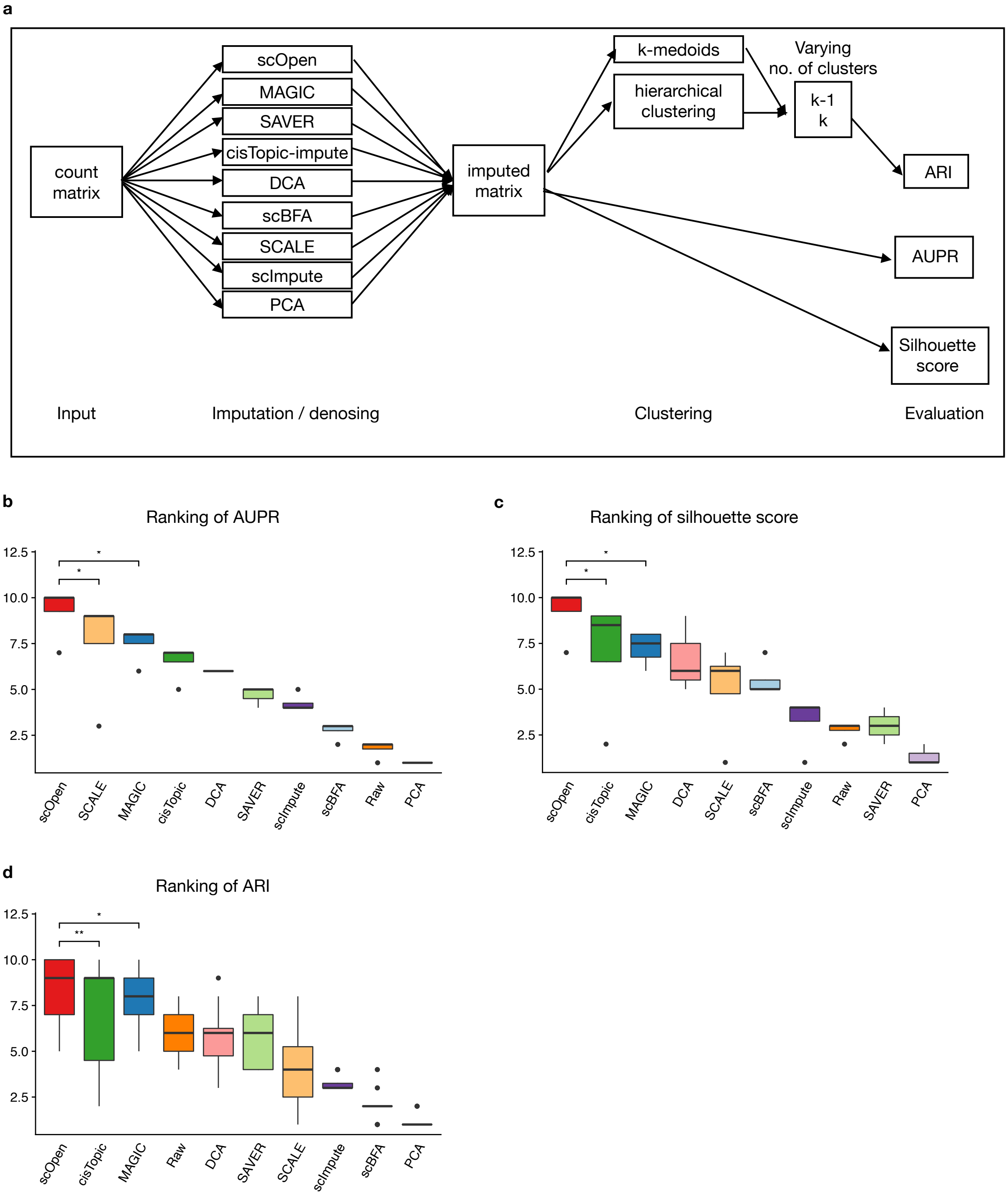


**Supplementary Fig. 2. Evaluation of imputation methods. a,** Experimental design for benchmarking competing methods. **b,** Ranking of imputation methods in terms of average AUPR for each benchmarking dataset. Methods are ordered by median value of ranks. Wilcoxon Rank Sum test was used to compare scOpen with SCALE and MAGIC. The asterisk means that the method is outperformed by scOpen with significance level of 0.5. **c,** Ranking of imputation methods using silhouette score as metric for each dataset. **d,** Ranking of methods using ARI as metric for benchmarking datasets. Asterisks denote statistical significance: * $p < 0.05$, *** $p < 0.001$

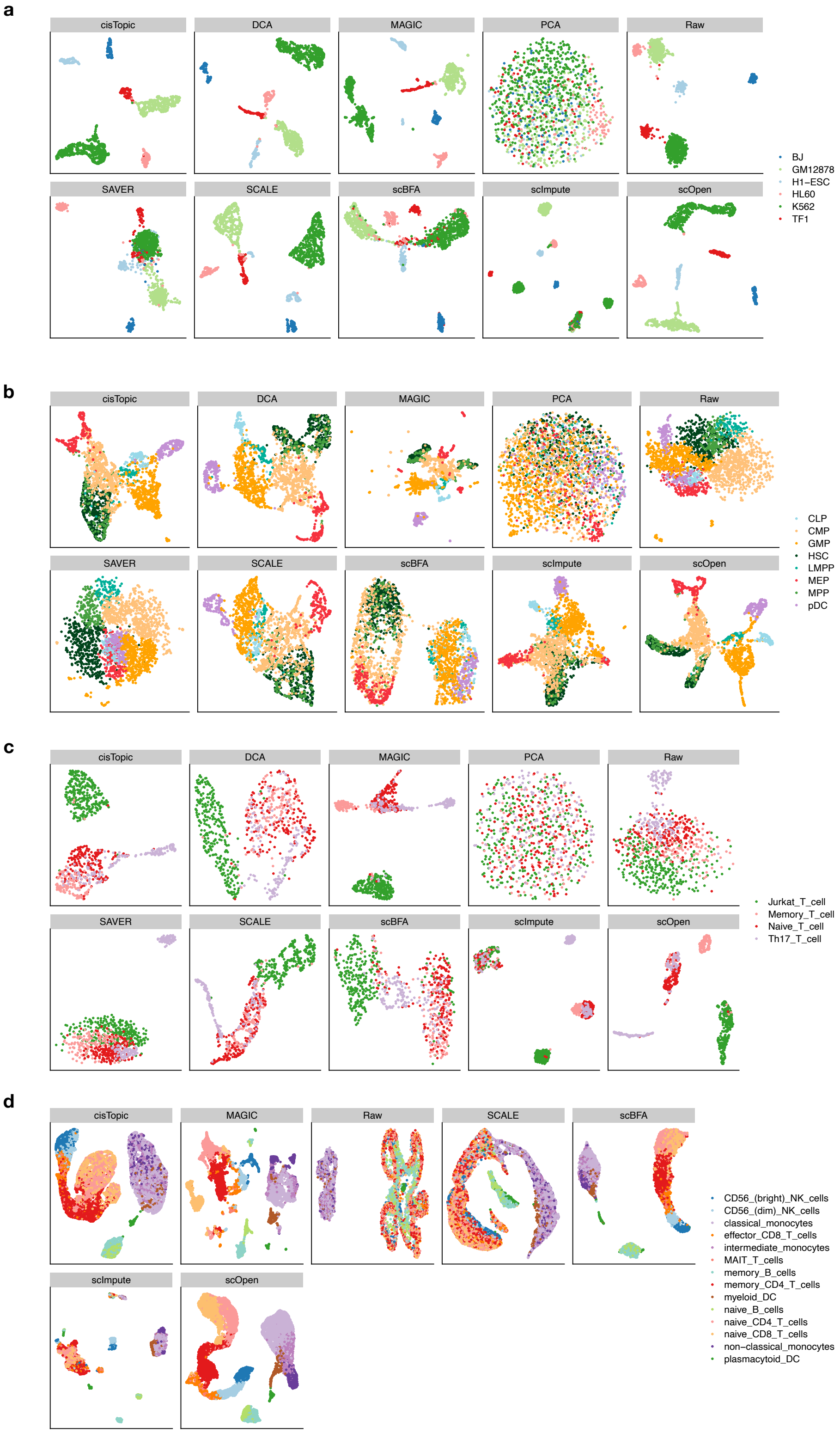


**Supplementary Fig. 3. Visualisation of imputation methods on benchmarking datasets. a,** UMAP embedding of scOpen, cisTopic, DCA, MAGIC, SAVER, scImpute, PCA and the raw data for cell line dataset. **b,** Same as **a** for Haematopoiesis. **c,** Same as **a** for T-cells. **d,** Same as **a** for multi-omics PBMC.

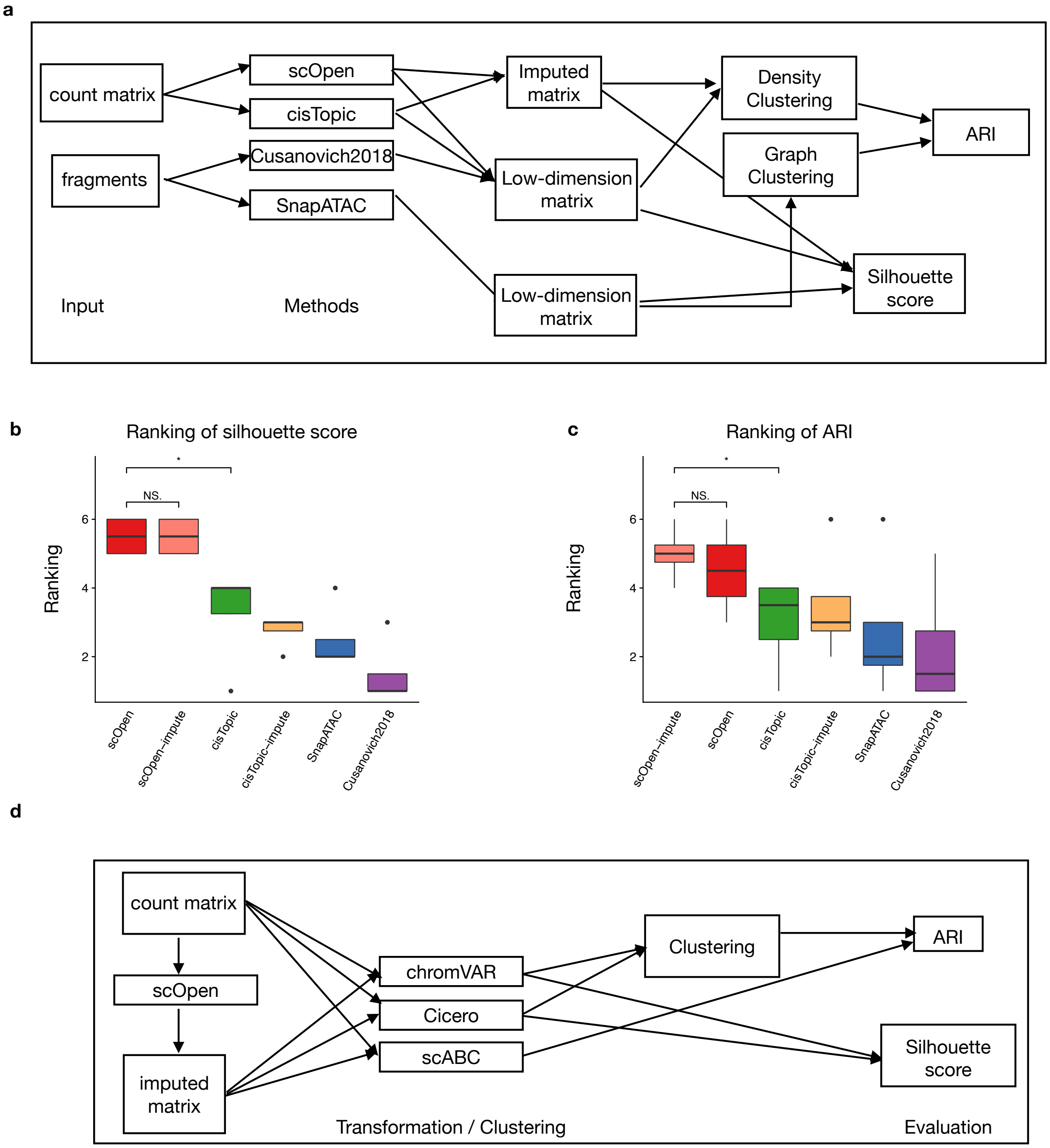


**Supplementary Fig. 4. Experimental design for benchmarking scOpen. a,** Experimental design for benchmarking of clustering pipelines for scATAC-seq. **b,** Boxplot showing the rank of silhouette score for clustering pipelines. We compared the top ranked method (scOpen) with the top-2 runner-up methods using Wilcoxon Rank Sum test. Asterisk denote statistical significance: * $p < 0.05$. **c,** Same as **b** for ARI. **d,** Experimental design for benchmarking downstream analysis methods between using raw or scOpen estimated matrix as input. The results were evaluated in terms of clustering with ARI as metric and silhouette score. Clustering was performed using the built-in method in each method

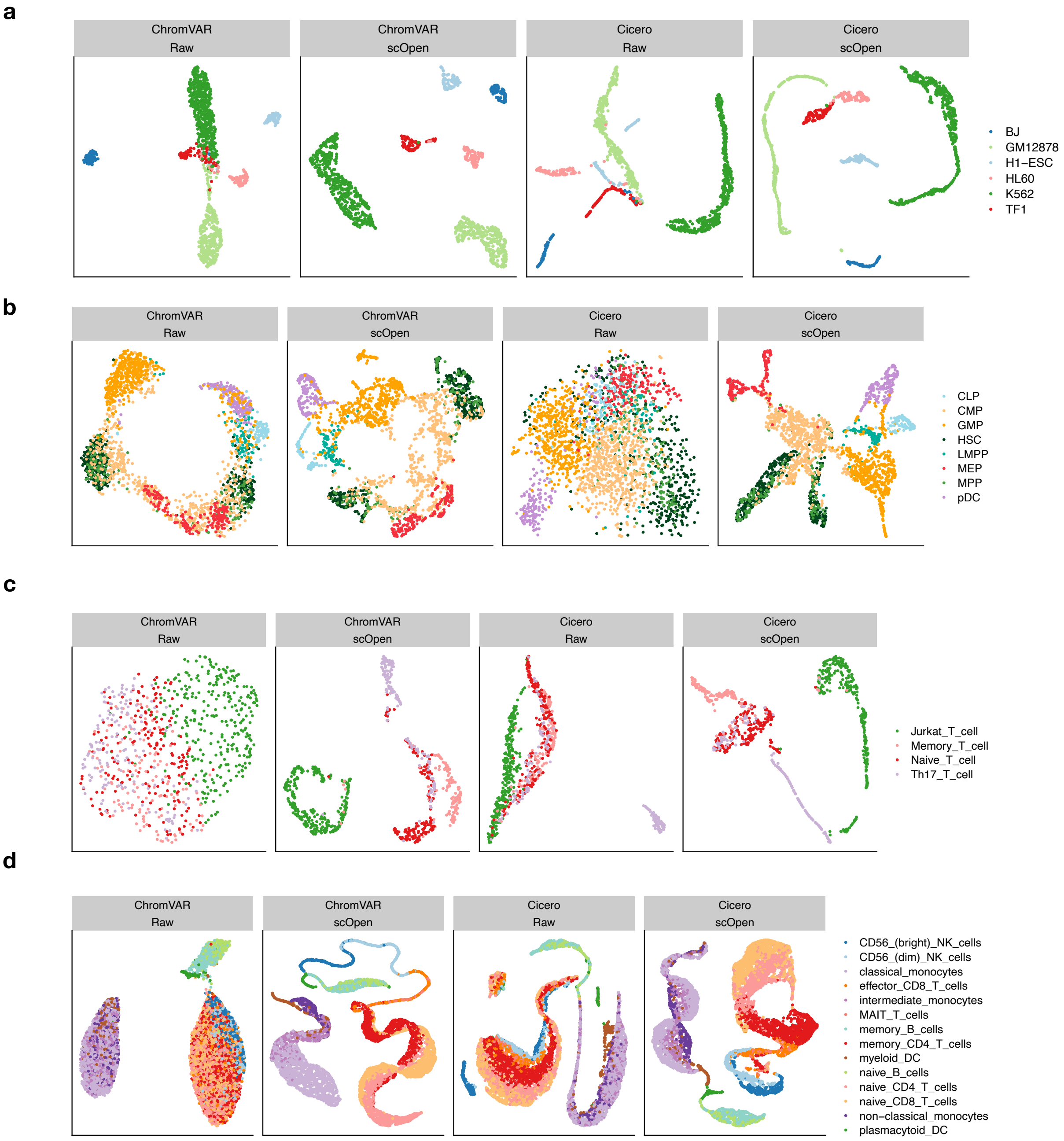


**Supplementary Fig. 5. Benchmarking and visualization for downstream analysis of scATAC-seq. a,** UMAP embedding of ChromVAR and Cicero transformed data using either raw or scOpen estimated matrix as input for cell line dataset. **b**, Same as **a** for Haematopoiesis. **c**, Same as **a** for T-cells. **d**, Same as **a** for multi-omics PBMC.

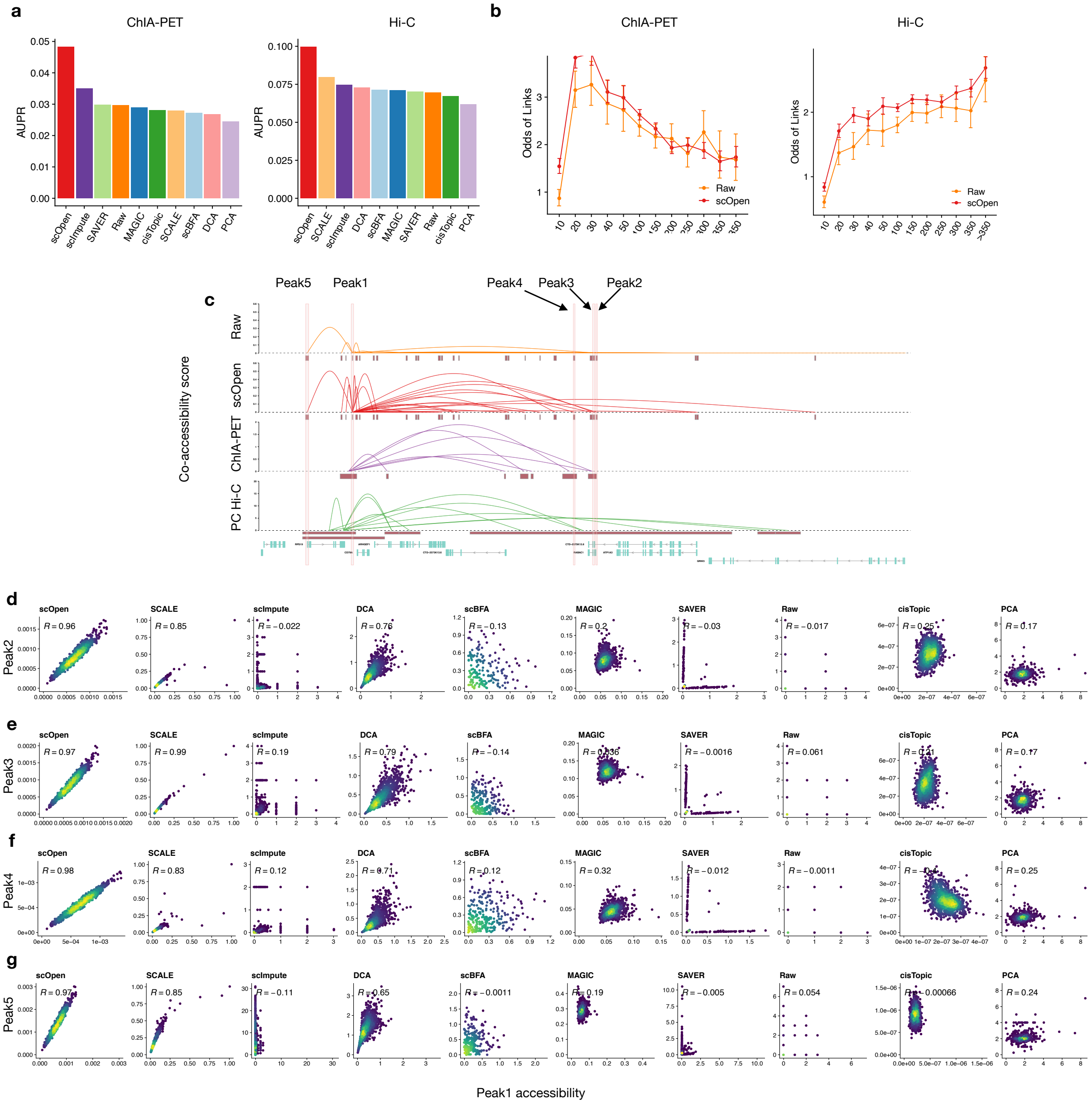


**Supplementary Fig. 6. Benchmarking of Cicero predicted links.** **a,** Barplots showing AUPR of the predicted peak-peak co-accessibility links using either raw or imputed matrix with H1-ESC single cell ATAC-seq data. Analysis was executed with Cicero. Left, links are evaluated using ChIA-PET data as true labels. Right, links are evaluated using Hi-C data as true labels. **b,** Odds ratio ($y$-axis) of Cicero predicted co-accessible sites (n = 3853260) also supported by pol-II ChIA-PET (left) and PC Hi-C (right) vs. distance between sites ($x$-axis). Error bars indicate 95% confidence intervals calculated using Fisher's exact test. Odds ratio superior than 1 indicates a positive relationship. **c,** Visualisation of co-accessibility scores (y-axis) of Cicero predicted with raw and scOpen estimated matrices contrasted with scores based on RNA pol-II ChIA-PET (purple) and promoter capture Hi-C (green) around the *CD79A* locus (x-axis). We highlight 4 links supported by Hi-C in this locus. **d,** Scatter plots showing single cell accessibility scores from raw data or estimated by imputation methods for peak1-to-peak2 link as shown in **c**. **e,** Same as **d** for link peak1-to-peak3. **f,** Same as **d** for link peak1-to-peak4. **g,** Same as **d** for link peak1-to-peak4.

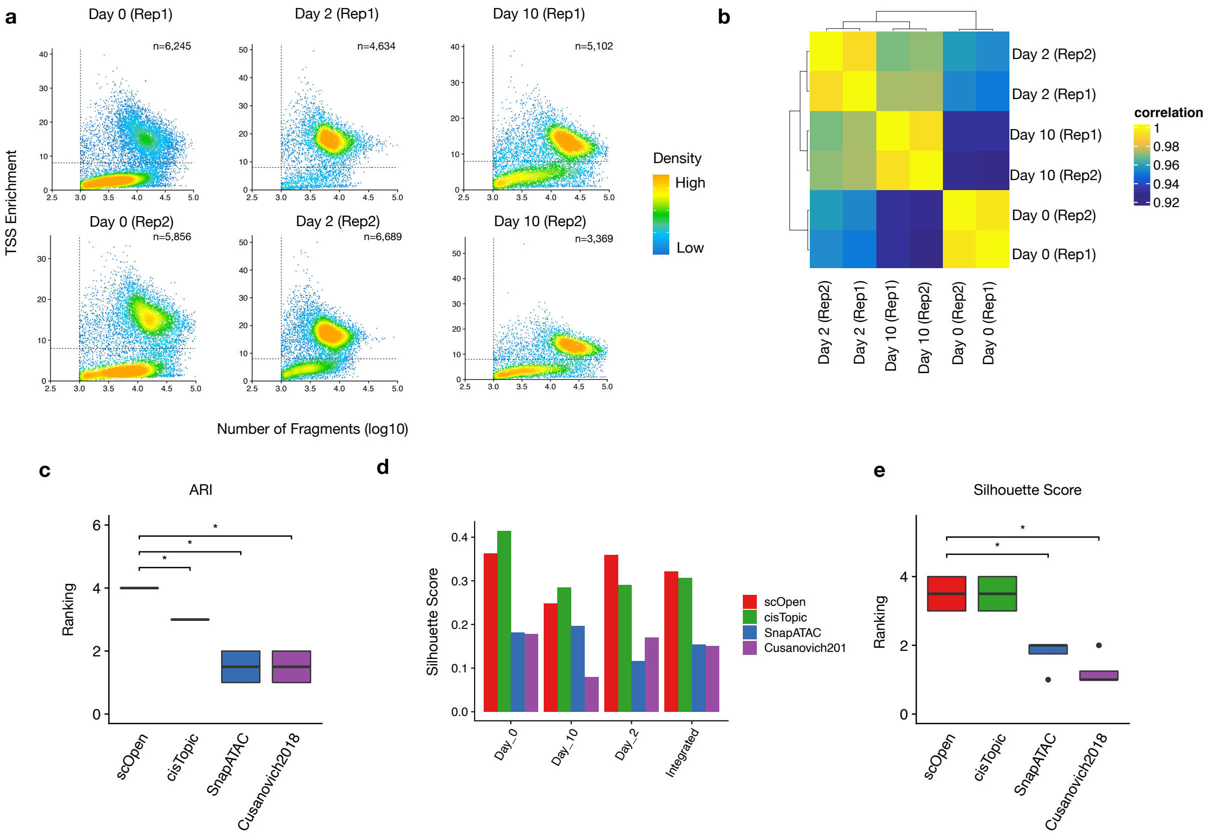


**Supplementary Fig. 7. Quality control and benchmarking of UUO scATAC-seq a,** Scatter plot showing number of unique fragments vs. TSS enrichment of UUO scATAC-seq for each sample. Each dot represents a cell and the dash lines represent cut-off used for cell filtering. The number of cells that pass filtering is shown on right-upper corner. **b,** Heatmap showing the correlation between samples. **c,** Rank of ARI for distinct scATAC-seq dimension reduction/clustering pipelines. Wilcoxon Rank Sum test was used to compare scOpen with SnapATAC and Cusanovich2018. Asterisk denote statistical significance: * $p < 0.05$. **d,** Barplot showing silhouette score of each sample by using distinct dimension reduction/clustering pipelines after data integration and label transferring. **e,** Same as **c** for silhouette score.

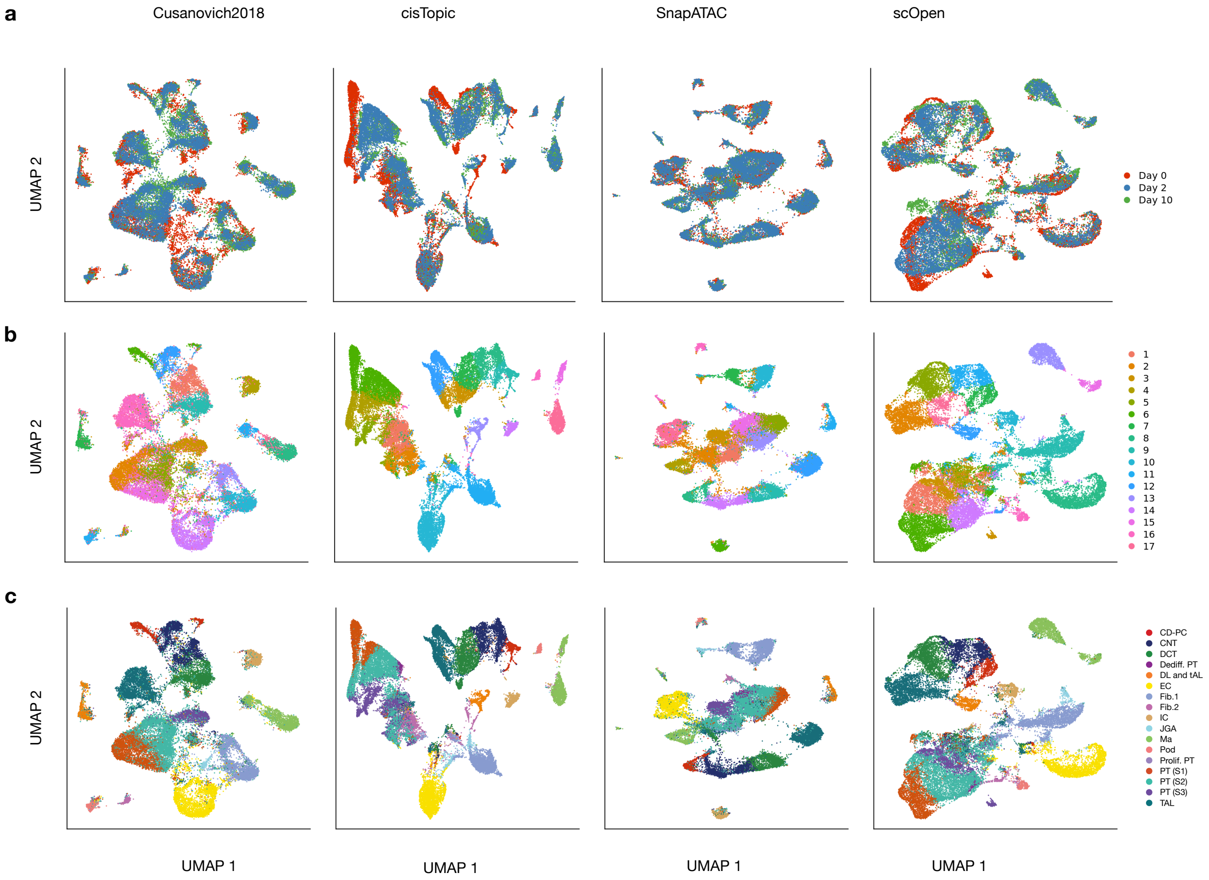


**Supplementary Fig. 8. Visualization of UUO scATAC-seq a**, UMAP plots showing data integration results by using different scATAC-seq dimension reduction/clustering pipelines. Each dot represent a cell and cells are coloured by time point. **b,** UMAP plots showing clustering results for each dimension reduction method. Cells are coloured by cluster. **c** UMAP plots showing label transfered results from a public UUO scRNA-seq dataset. Cells are coloured by predicted label.

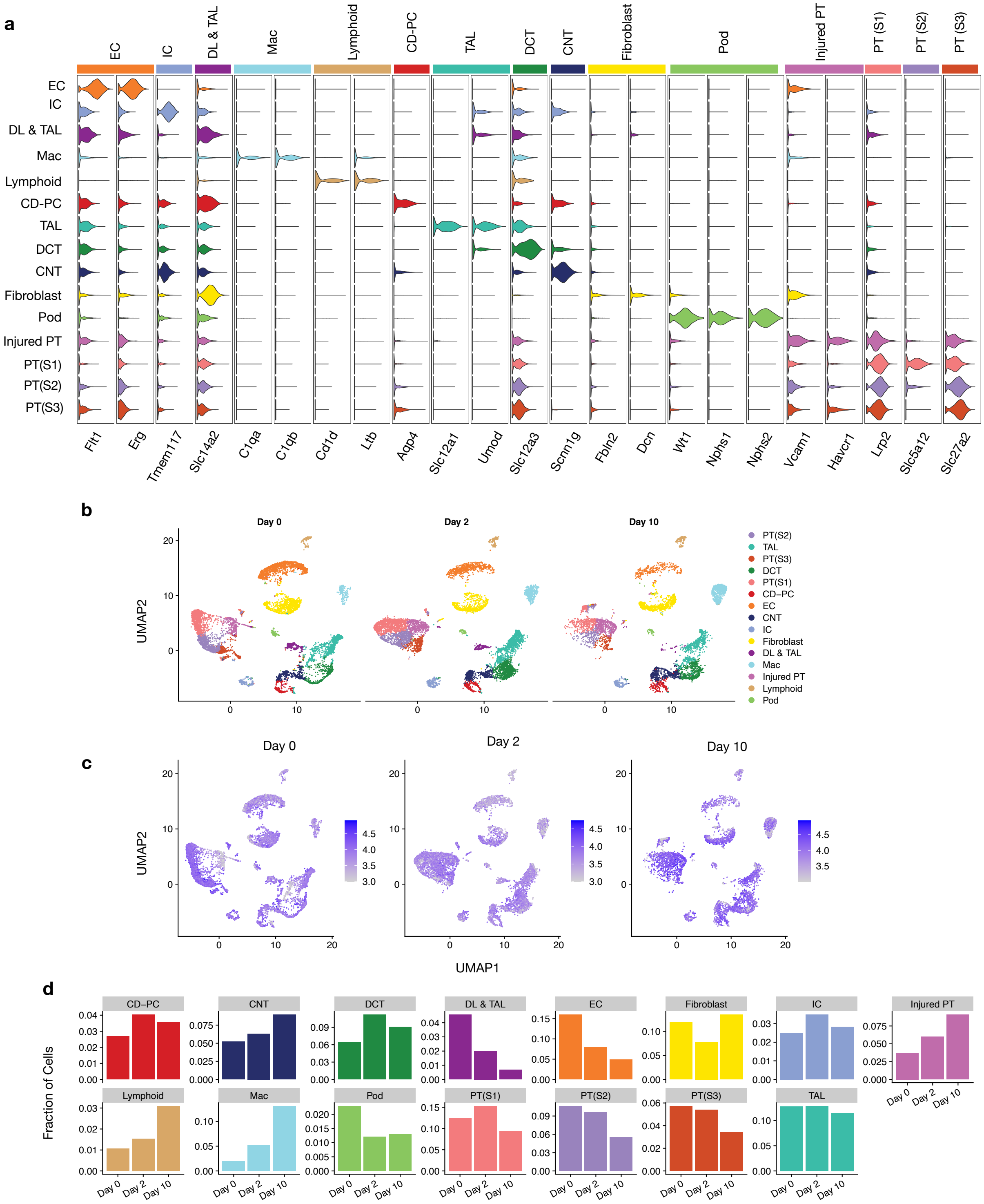


**Supplementary Fig. 9. Visualization of UUO annotation a,** Violin plot showing cluster-specific (y-axis) gene accessibility score associated to known marker genes for kidney cells (x-axis). **b,** Scatter plots showing condition-specific UMAP visualization of UUO scATAC-seq data for each day. **c,** Visualisation of data quality for **b**. Colours refer to number of fragments per cell (log10). **d,** Bar plots showing proportion of each cell type across different days. For abbreviation of cell types listed see legend of Fig. 3b.

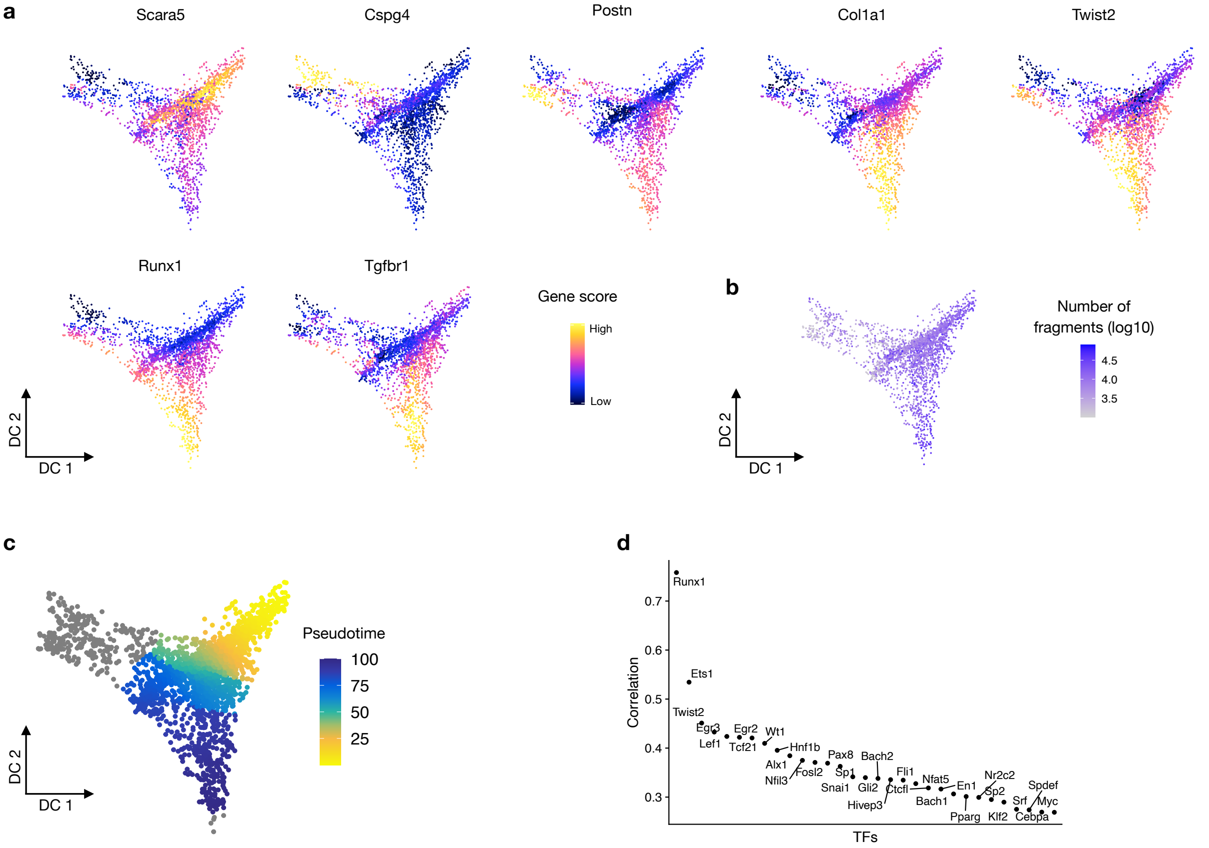


**Supplementary Fig. 10. Identification of TFs driving the differentiation of fibroblasts to myofibroblasts a,** Diffusion map embedding of fibroblast showing gene activity score for Scara5 (fibroblast), Cspg4 (pericytes), Postn (myofibroblasts), Col1a1 (myofibroblasts), Twist2, Runx1 and Tgfbr1. **b,** Visualisation of data quality for **a**. Colours refer to number of fragments (log10) per cell. **c,** Visualization of trajectory from fibroblast to myofibroblasts. Colour refer to pseudo-time during of the trajectory. **d,** Correlation of gene activity score and motif deviation (associated to **Fig. 4c**) along the trajectory for selected TFs. Each dot represent a TF and TFs are sorted by their correlation. Runx1 has the highest correlation, followed by ETS1 and Twist2. Of these, Runx1 and Twist2 have an increase in gene scores and TF activity along pseudo-time.

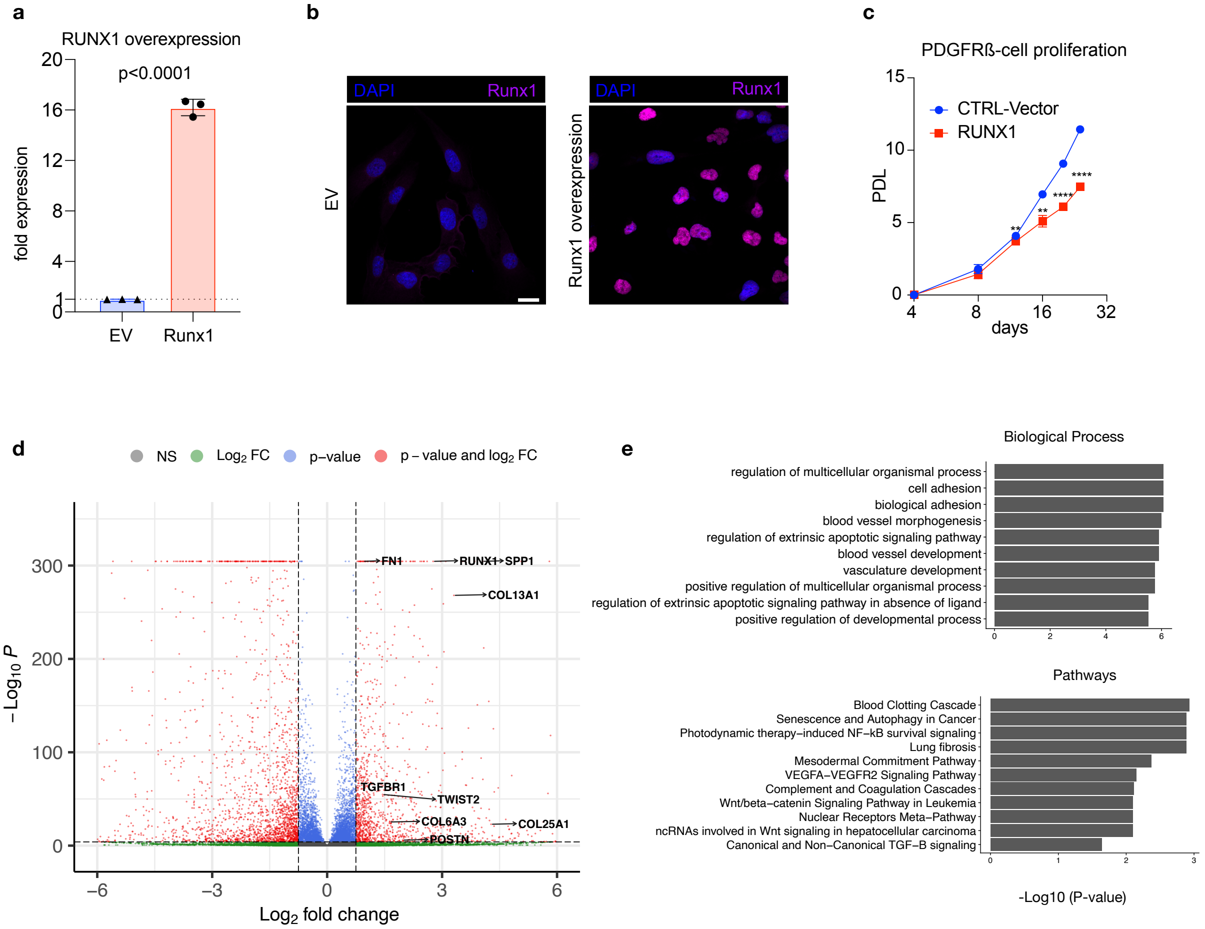


**Supplementary Fig. 11. Validation of Runx1 a,** Expression of Runx1 by qPCR after lentiviral Runx1 overexpression in a human kidney PDGFRb+ cell line (n=3). **b,** Immunofluorescence (IF) staining of Runx1 in human kidney PDGFRb+ cells lentivirally transduced with either empty vector (EV) or an Runx1 overexpression construct (Runx1). **c,** Population doubling of Runx1 overexpressing cells vs. control (EV) (n=3). **d,** Volcano plot showing differential expression analysis between Runx1 overexpression vs. control. Each dot represents a gene and dashed lines represent the thresholds (x = 1.5 and y = 5) used for selection of DE genes. Colours refer to significance given different criteria. **e.** Barplot showing GO enrichment using up-regulated genes from Runx1 overexpression. Top 10 terms are shown for Biological Process and WikiPathways.

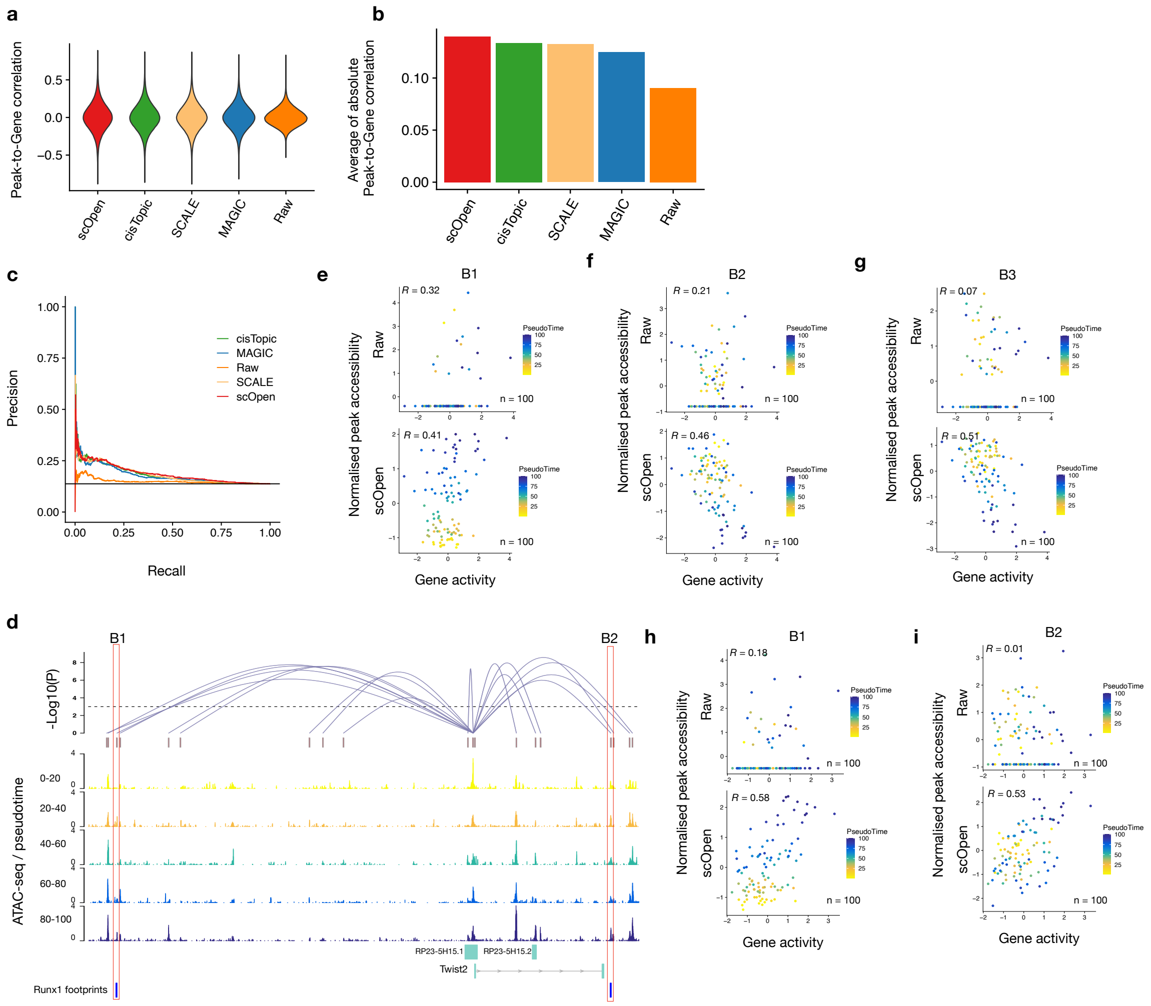


**Supplementary Fig. 12. Visualization and validation of Runx1 target genes a,** Violin plot showing predicted Peak-to-Gene (P2G) links using raw or imputed matrix from scOpen, cisTopic, SCALE and MAGIC. Methods are sorted by average absolute value. These methods were selected as top performing methods (see Sup. Fig. 2b). **b,** Average absolute value of correlation of P2G links. **c,** AUPR of predicted Runx1 target genes using from raw, scOpen, SCALE or cisTopic matrix. Related to **Fig. 4h**. **d,** Peak-to-Gene links of Twist2 in fibroblast cells. Each loop refers to a link and height of loop represent the significance (dash line represents threshold of FDR= 0.001). ATAC-seq tracks were generated from pseudo-bulk profiles of cell along the pseudo-time. Peaks with binding sites of Runx1 supported by ATAC-seq footprints are highlighted (B1-B2). **e,** Scatter plot showing gene activity of Tgfbr1 and linked peak accessibility from raw (upper) or scOpen estimated matrix (lower) for peaks B1 indicated in **Fig. 4i** of the main manuscript. Each dot represents cells in a given pseudo-time and the colour refers to pseudo-time. The correlation is shown on left-upper corner. Both gene activity score and peak accessibility are z-score normalised. **f,** Same as **e** for Peak-to-Gene link B2. **g,** Same as **e** for Peak-to-Gene link B3. **h,** Scatter plot showing gene activity of Twist2 and B1 peak accessibility from raw (upper) or scOpen estimated matrix (lower). **i,** Same as **f** for the Peak-to-Gene B2.

**Supplementary Tab. 1. Statistics of data sets used in this study**. The number of detected cells, number of regions (peaks), fraction of non-zero entries, average number of reads per cell, fraction of reads in peaks (FRIP) and total number of valid reads are shown below. For comparison purposes, we also included similar statistics for a scRNA-seq data corresponding to the Hematopoiesis scATAC-seq data. We observed a higher sparsity in scATAC-seq (0.039 vs 0.119) despite the fact scATAC-seq data has 7 times more reads per cell than scRNA-seq.

| Dataset | Type | Cells | Features | Non-zeros | Reads per cell | FRIP | Reads | Optimal Rank |
|---|---|---|---|---|---|---|---|---|
| Cell lines | scATAC-seq | 1,224 | 125,647 | 0.036 | 41,467.80 | 0.248 | 50,756,587 | 8 |
| T cells | scATAC-seq | 765 | 49,344 | 0,033 | 14,963.39 | 0.418 | 11,446,993 | 8 |
| Hematopoiesis | scATAC-seq | 2,210 | 109,418 | 0.039 | 34.656.15 | 0.272 | 76,590,091 | 9 |
| Hematopoiesis | scRNA-seq | 14,432 | 12,558 | 0.119 | 5.209,45 | NA | 75,182,840 | NA |
| PBMC | scATAC-seq | 10,032 | 106,935 | 0.067 | 13,486 | 0.714 | 457,001,034 | 7 |
| UUO | scATAC-seq | 31,129 | 150,593 | 0.042 | 13,933 | 0.467 | 419,794,555 | 18 |
